# A pipeline for systematic comparison of model levels and parameter inference settings applied to negative feedback gene regulation

**DOI:** 10.1101/2021.05.16.444348

**Authors:** Adrien Coulier, Prashant Singh, Marc Sturrock, Andreas Hellander

## Abstract

Quantitative stochastic models of gene regulatory networks are important tools for studying cellular regulation. Such models can be formulated at many different levels of fidelity. A practical challenge is to determine what model fidelity to use in order to get accurate and representative results. The choice is important, because models of successively higher fidelity come at a rapidly increasing computational cost. In some situations, the level of detail is clearly motivated by the question under study. In many situations however, many model options could qualitatively agree with available data, depending on the amount of data and the nature of the observations. Here, an important distinction is whether we are interested in inferring the true (but unknown) physical parameters of the model or if it is sufficient to be able to capture and explain available data. The situation becomes complicated from a computational perspective because inference and model selection need to be approximate. Most often it is based on likelihood-free Approximate Bayesian Computation (ABC) and here determining which summary statistics to use, as well as how much data is needed to reach the desired level of accuracy, are difficult tasks. Ultimately, all of these aspects - the model fidelity, the available data, and the numerical choices for inference and model selection - interplay in a complex manner. In this paper we develop a computational pipeline designed to systematically evaluate inference accuracy for a wide range of true known parameters. We then use it to explore inference settings for negative feedback gene regulation. In particular, we compare a spatial stochastic model, a coarse-grained multiscale model, and a simple well-mixed model for several data-scenarios and for multiple numerical options for parameter inference. Practically speaking, this pipeline can be used as a preliminary step to guide modelers prior to gathering experimental data. By training Gaussian processes to approximate the distance metric, we are able to significantly reduce the computational cost of running the pipeline.

## 1 Introduction

Mathematical modeling is an important tool to study gene regulatory networks (GRNs) in single cells. These models exist on many levels of fidelity, ranging from deterministic Ordinary Differential Equations (ODE) to discrete stochastic well-mixed models, to detailed spatial stochastic models. In practice, the choice of a mathematical model has a subjective element to it and the choice depends both on the question under study, the available data, and the computational budget.

There have been many studies where modelers have simulated models at different levels of fidelity to study the same biological system. Often these studies have had the same aim of capturing either qualitative properties or relatively coarse-grained summary statistics, both of which could in principle be captured by any of the models. One commonly studied biological system is the Hes1 negative feedback system. This system produces oscillatory dynamics which are thought to provide a clock for the segmentation of the somites during embryogenesis [25]. This system has spawned models of different levels of biological fidelity, including models comprised of ordinary differential equations [6], delay differential equations [40, 28], partial differential equations (PDE) [10], stochastic differential equations [24] and spatial stochastic simulations [52]. These models were used to simulate the Hes1 system with the aim of producing oscillatory dynamics of the Hes1 protein and messenger RNA (mRNA) with a period of approximately 2 hours. All models were able to produce the desired oscillatory behaviour, but in the ODE case an extra intermediate reaction had to be introduced. This deployment of a variety of modelling approaches is not unique to the Hes1 intracellular pathway, the p53 signalling pathway [15, 53] and NF-*κ*B pathway [57] have similar variety in the fidelity of models produced to capture simple summary statistics. It is not clear in these studies whether the data alone necessitated a higher fidelity model. Outside of the intracellular modelling space, there have been many similar studies in the cancer modelling space that have used models of differing fidelity to capture tumour growth time series data. These models have often been either ODE or PDEs or even agent based models with huge differences in computational expense and model complexity [16]. Again though, it is not clear if and when the available data clearly motivated one approach over another. A central issue is whether the models are developed with the goal to infer parameter values from experimental data or if they are developed to predict some future state of the system. In the latter case, some have argued that the precise parameter values may be of less concern and may even take biologically infeasible values so long as the model is a good predictor of the system [22].

In some situations the question at hand clearly motivates a spatial model over a well-mixed one, such as if we were to study the effect of location and numbers of membrane receptors on downstream signaling in a signaling cascade. However, in situations where the question could in principle be addressed using models of various different levels of fidelity, and when the choice is driven by observed data, the situation is less clear. In particular, it is common that quantitative experimental observations are “well-mixed” in nature, such as time series data of total protein or mRNA counts in cells, or distribution data from large cell populations from e.g., fluorescent activated cell sorting (FACS). An interesting question then is in which situations it is motivated to use a high-fidelity spatial model even though the observations are more coarse-grained in nature. This question can be expected to depend critically on what the goal of the modeling is, for example, is the goal simply to capture the qualitative trends in data, or is the goal to identify model parameters that are in good quantitative agreement with the true biochemical and physical parameters, such as diffusion constants and kinetics rate constants? This is fundamentally a hard problem to address due to the lack of ground truth (both for the model and for the parameter). But if enough computational power is available, we can scan through the space of possible parameters and evaluate how inference would perform were the postulated parameters the ground truth (synthetic data). While the true posterior distribution is out of reach, we can still analyze how the estimated posterior relates to the true parameters for various choices of models and for various types of observations. In this paper, we develop such a computational pipeline in which we generate synthetic ground truth data using a high-fidelity spatial model (simulated using Smoldyn [4]) for a wide range of possible true parameters (controlling the degree of diffusion control of the system), then systematically compare inference tasks for the spatial model, the well-mixed model and a coarse-grained multiscale model.

However, since it is necessary to use likelihood-free, or simulation-based approximate inference, there are several numerical considerations for accurate inference, apart from the question of the simulator and how much data is available. In particular, ABC, the mosty widely used method, relies critically on the chosen distance metric and summary statistics. In the end, both the model fidelity, the data, and the numerical choices for inference need to be studied simultaneously. The fact that ABC requires a large number of potentially expensive simulations becomes a practical hinder to such large-scale studies. Here we train Gaussian Processes (GPs) to approximate the distance metric, and are in this way able to significantly reduce the computational cost of running the pipeline.

While ideally models would be constrained using sufficiently rich experimental data in practice there are often limitations to the amount and kind of data available. The modeling of cellular functions requires sensitive measurement of various molecular species such as mRNA and proteins. For data captured at the intracellular level, there are often trade offs between the richness of the time series, the number of replicates and the level of spatial information captured. Traditionally used population-averaged techniques like Western blots, Northern blots and enzyme-linked immunosorbent assay (ELISA) do not capture the important details at the single cell level. More modern techniques such as mass spectrometry (MS) generally lack the sensitivity to detect the small amounts of proteins present in individual cells [1, 45]; however, recent developments in MS have made progress towards uncovering single-cell proteomes [8]. Flow cytometry and mass cytometry (e.g., CyTOF) can detect proteins in single cells, but developing sample standards for quantification has proved challenging [5]. In very recent years, the digital proximity ligation assay (dPLA) was developed. dPLA provides the ability to take direct digital measurements of protein and mRNA copy numbers in single mammalian cells [3]. In dPLA, digital PCR (dPCR) was used to quantify proteins detected with a pair of oligonucleotide-tagged antibodies called PLA probes. Previously published PLA methods enabled multiplexed simultaneous protein and mRNA measurements from single cells. It was noted though that their quantitative polymerase chain reaction (qPCR) readout limits the sensitivity of the measurements [12]. The use of the dPCR readout provides significantly improved resolution and limits of detection [56], which allows direct quantification of protein copy numbers in individual mammalian cells. This advance, along with the use of dPCR for mRNA quantification allows for simultaneous measurement of both mRNA and protein albeit at low time resolution (with readings captured every 10 minutes) [36]. All of these experimental technologies come with different costs and can require different levels of experience to gather, hence it is important to know when enough data is captured to warrant the use of a more sophisticated model. To that end, in this study we use a synthetic data set that mimics the cutting edge of what is possible experimentally, i.e. simultaneously capturing mRNA and protein copy number data at the single cell level for various cells and time points, to address the question of how much data is needed to warrant a higher fidelity model.

There are a growing number of studies investigating parameter inference in the presence of different kinds of data. In [24], it was demonstrated that MCMC methods for stochastic differential equations provide practical algorithms for estimating the parameters of simple dynamic regulatory and signaling systems even when the time series data are coarse. Furthermore, it was reported that if one had access to good quality temporally resolved data, one could also obtain information about stochastic modeling parameters and population sizes. In [34], Kursawe *et al.* investigated the performance of parameter inference using a vertex model for cell mechanics and image data. They showed that estimating the noise by having several realizations of the observed data was critical for reliable inference. Harrison *et al.* [23] quantified the effect of noise and data density on the posterior estimate and compared ABC to the particle Markov Chain Monte Carlo method (pMCMC). They showed that, when applicable, pMCMC performed better, although ABC was more general and more easily parallelizable.

Some doubts have been raised regarding the validity of the posterior distributions generated with ABC [44, 54]. Specifically, and although they are fundamental requirements for the well-posedness of the inverse problem, identifiability [38] and sufficiency [43] may not be attainable in practice. Yet, this need not be the end of the story. Firstly because identifiability *can* be demonstrated for some simpler models [7], and secondly because insights can still be gained from models where the true solution is known [37].

In this paper, we show that such an approach is indeed feasible for ABC. We propose a computational pipeline to evaluate the accuracy of ABC in different scenarios. Specifically we evaluate the performance of ABC throughout the parameter space while keeping the cost down using Gaussian processes to approximate the distance metric. We then analyze the accuracy of the resulting posterior distribution with respect to the true parameters. We can then repeat this procedure for different models, summary statistics or even data sets (e.g., when measuring proteins or when measuring mRNA) and determine which setup gives the best performance. This preliminary analysis can then be used to guide practitioners to choose models and design experiments. Contrary to other analytical approaches to guide modelers and optimize experiments [55, 18], our approach is purely computational and does not require the ability to derive analytical formulae from the model formulation. Hence, models of arbitrarily high complexity can be used. With this approach we can answer questions such as:

- How much data is needed to reach a given level of inference accuracy?
- What features are worth measuring in an experiment if the goal is to identify parameters?
- Which modeling level is appropriate?
- Which summary statistics or distance metric should be used?

We exemplify this procedure through different scenarios based on a canonical negative feedback gene regulation network motif for three models of various fidelities. The remainder of this paper is organized as follows: in Section 2, we give some background on stochastic modeling and likelihood-free Bayesian inference. In Section 3, we explain our pipeline in detail. In Section 4 we illustrate how this pipeline can be used in four different scenarios. Finally in Section 5 we discuss our contributions and future areas of research.

## 2 Background

### 2.1 Stochastic models for chemical kinetics

Stochastic chemical kinetics in single cells can be modeled at various levels, from stochastic differential equation to detailed particle models [21]. Yet, the choice of modeling level is not always easy to do *a priori*. In particular, models including spatial details about the distribution of molecules throughout the cells can reveal new insights but come with a significant increase in computational cost [9].

A popular modeling framework is the Chemical Master Equation (CME) [19]. In the CME formalism, the system is represented by a state vector **x** where each row represent the molecular count of a given species. The probability distribution for a system of *n* species and *m* reactions is given by the solution of the master equation:

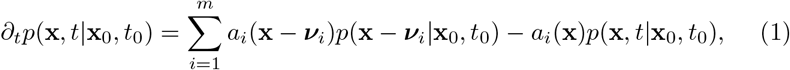

where **x** is the state vector of the system, *a_i_*(**x**) and ***ν****_i_* are the propensity and the stochiometric vector of reaction *i*, respectively, and *x*_0_ is the state of the system at time *t*_0_.

Unfortunately, solving the CME numerically is in most practical cases intractable. It is however possible to generate realizations of the CME using Gillespie’s Stochastic Simulation Algorithm (SSA) [20]. One fundamental assumption of the CME is that molecules inside the cell are *well-mixed*, i.e. there is enough time between reactions for the molecules to diffuse uniformly across the cell. In other words, the CME does not include spatial details about the location of each molecules.

In [11], we presented a technique to include some degrees of spatial details into the CME framework. By dividing the cell into compartments and computing transition rates between these compartments, we are able to include some spatial information and approximate more detailed models for only a marginal increase in computational cost.

Another, more standard generalization of the CME to include spatial details is the Reaction-Diffusion Master Equation (RDME) [14, 51]. In the RDME framework, space is discretized into small voxels. Every voxel is assumed to be well mixed and reaction can only occur between molecules belonging to the same voxel. Additionally, molecules can diffuse to neighboring voxels, depending on the geometry of the discretization and of the diffusion rate.

Other, even more detailed methods track the position of each molecule in continuous space. For instance, in the Smoluchowski diffusion limited model, molecules diffuse in space following Fick’s second law:

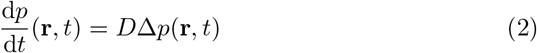

where **r** is the position of a molecule, *D* its diffusion constant and *p* the probability distribution of its position. Molecules are then modeled as hard spheres that react upon collision with a given probability. Solving the Smoluchowski equation in the general case is an open problem. One approach to circumvent this issue is to discretize time and rely on approximations to determine when two molecules collide and potentially trigger a chemical reaction. This approach is used in e.g. Smoldyn [4]. Another approach consists in isolating pairs of molecules or single molecules in protective domains where the Smoluchowski equations can be solved analytically using Green’s Function Reaction Dynamics. This is the approach used in e.g. eGFRD [50].

There is a well defined mathematical hierarchy between these approaches [21, 49]. As a matter of fact, it is known that in the limit of infinite diffusion, spatial approaches will converge towards the CME. However, in practice, it is difficult to determine if diffusion is “fast enough” for the CME to be a valid approximation of the chemical system under scrutiny. Using a more detailed model will be more accurate, but at the price of a higher computational cost. Balancing these two aspects is a critical question in modeling.

In a previous study [11], we considered three stochastic models including various level of details and compared how they related to each other in terms of accuracy in the context of the Hes1 system presented in [52]. In Section 4 we will use the same models to illustrate how our pipeline can used to compare these models in a Bayesian parameter inference setting.

### 2.2 Likelihood-free Bayesian Inference

Given a mathematical model **y** = *f* (*θ*), the goal of parameter inference is to fit *f* to observed data **y***_o_*, i.e., estimate the parameters *θ_o_* that give rise to simulated data **y***_sim_ = f* (*θ_o_*) such that **y***_sim_* = **y***_o_*. As the model *f* is stochastic, in reality the equality condition is too strict and can never be fulfilled exactly. Typically the equality condition is replaced by a relaxed form involving a threshold. Also, we note that for all but very simple models of academic interest, the likelihood function corresponding to *f* is unavailable. Therefore, parameter estimation must proceed in a likelihood-free manner making use of the observed data, and access to the simulation model *f*.

The most popular family of parameter estimation methods in the likelihood-free setting is approximate Bayesian computation (ABC) [48]. ABC parameter inference begins with the specification of a *prior* distribution *p*(*θ*) over the parameters *θ*, representing the parameter search space. The ABC rejection sampling algorithm then samples *θ*_sim_ ~ *p*(*θ*), and simulates **y**_sim_ = *f* (*θ*_sim_). The simulated time series **y**_sim_ must now be compared to **y***_o_* to validate whether the two time series were sufficiently close. If so, then *θ*_sim_ is deemed to be *accepted*, else *rejected*. This sample-simulate-compare rejection sampling cycle is repeated until enough amount of accepted samples are obtained. The set of accepted samples then form the estimated posterior distribution *p*(*θ*|**y***_o_*), solving the parameter inference problem.

The comparison between time series’ **y***_o_* and **y**_sim_ is typically performed in terms of *k* low-dimensional *summary statistics* **S** = *S*_1_(**y**), …, *S_k_*(**y**) or features of the time series (e.g., statistical moments). This is due to the *curse of dimensionality* when comparing rich high-dimensional time series (detailed discussion in Chapter 5 of [48]).

Summary statistic selection is a well-studied problem, and there exist methods to select *k* informative statistics out of a candidate pool of *m* total statistics. A thorough treatment of the topic can be found in [43, 48]. A well motivated method of selecting summary statistics is based on the notion of approximate sufficiency (AS) [31]. Summary statistics **S** are sufficient if adding a statistic *S*_new_ to **S** does not change the approximated posterior *p*(*θ*|**y***_o_*). The AS algorithm initiates tests for different statistics in random order [41], therefore in this work we will repeat summary statistic selection several times to compute the frequency of selection of each statistic. The most frequently selected statistics will be used in the parameter inference process. The candidate pool of statistics to choose from include the following statistics/features for each species.

|  |  |
| --- | --- |
| – sum of values | – kurtosis |
| – absolute energy | – longest strike below mean |
| – mean absolute change | – longest strike above mean |
| – mean change | – last location of maximum |
| – median | – first location of maximum |
| – mean | – last location of minimum |
| – length | – first location of minimum |
| – standard deviation | – maximum |
| – skewness | – minimum |

Therefore, in total there are 36 candidate statistics to choose from (18 for each species - mRNA and protein).

## 3 A Computational Pipeline for Systematic Parameter Inference Evaluation

Given that we want to be able to use models of high complexity it is usually not possible to rely on *a priori* mathematical analysis to determine the validity of ABC for a given setup. For example, information on system identifiability is often not available in practice, and numerical identifiability needs to be studied empirically. We can, however, systematically evaluate the performance of ABC using synthetic observed data sampled throughout the parameter space. Given the high computational cost, we use an approximation scheme based on Gaussian processes. This makes it possible to only generate the data once, prior to inferring parameters with ABC in various configurations.

### 3.1 Overview

We have developed a pipeline made up of two main parts, a data generation step and a parameter inference step. Figure 1 illustrates how these parts are combined. Starting with a prior distribution, we first simulate ground truth data for one parameter point using the highest model fidelity (Fig. 1A and then generate simulated data across the entire parameters range of the prior with the model meant for Bayesian inference (Fig. 1B). We then use this data to approximate the distance between observed (synthetic data generated with the highest fidelity) and simulated data. This approximate distance map (Fig. 1C) is then used to perform parameter inference without the need to simulate more data during this process (Fig. 1D).

**Figure 1:**
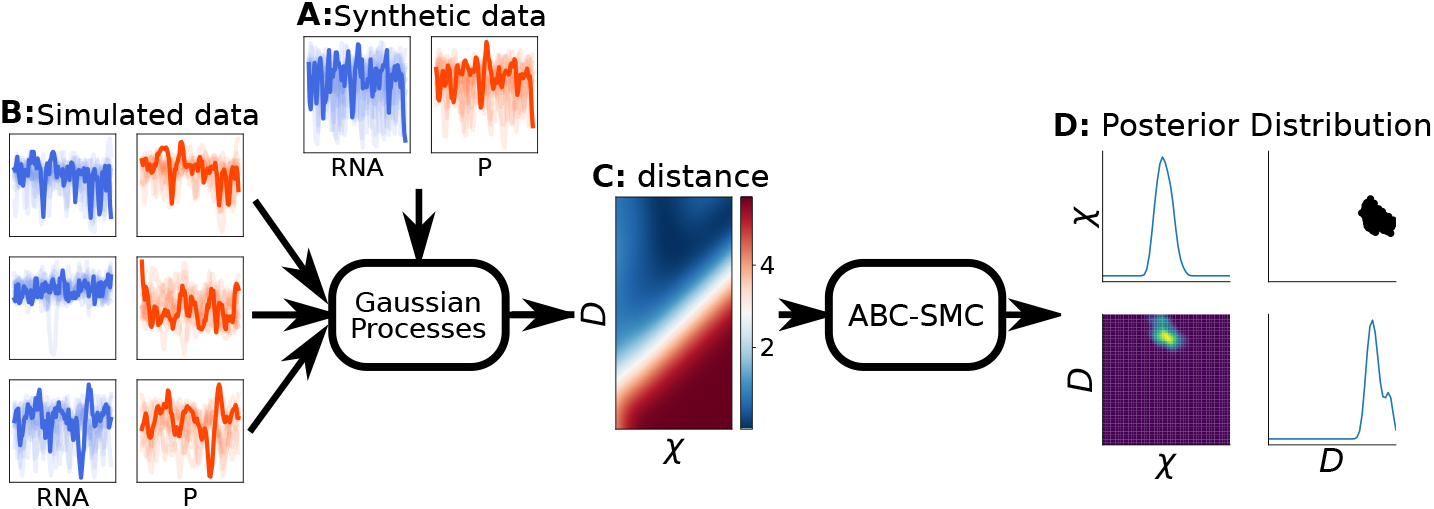
Parameter inference pipeline based on Gaussian processes. A distance metric based on one synthetic data set (A) is trained using all other simulated data sets (B). This distance metric (C) is then used in pyABC to infer the parameters of the synthetic data set using ABC-SMC (D).

This entire pipeline can then be executed multiple times using synthetic observed data sets from different regions of the parameter space. The core idea is to generate the data once and then reuse it first as synthetic observed data, and then as training data to approximate the distance function used in ABC. Thus, the same data set can be reused in various configurations, e.g. with different summary statistics or by subsampling the data in terms of number of trajectories or time samples. This in turn generates many posterior distributions which can be compared to the true parameters. By systematically measuring the discrepancy between posterior distribution and true known parameters, we can then build up error maps showing regions in parameter space where inference performs better or worse.

Analyzing the thus obtained error maps from various combinations of models, summary statistics and amounts of data can offer insights into which combination would be more or less likely to perform well when used against experimental data. We believe that this pipeline can be used as a pre-processing step to calibrate inference pipelines when the true parameters and model is unknown, assuming we believe that the model of the highest fidelity is fundamentally the best representation of reality of the simulators we consider. As we will show, the pipeline is in particular useful to reason about in which situations it is motivated to use which model fidelity in systems biology projects.

### 3.2 Gaussian Processes for Likelihood-Free Parameter Inference

ABC typically entails a slow convergence towards the estimated posterior, and may require several thousands of simulations to provide a reliable estimate. When simulations are expensive, and when we need to perform many such inference computations, this can represent a serious computational bottleneck. In this study, we set to make this cost as small as possible, so that we can repeat parameter inference experiments over wide prior ranges and many inference setups.

In [27], Järvenpää *et al.* describe how Gaussian processes can be used to approximate the distance function, or discrepancy, between simulated data and observed data and demonstrate how this technique can be used to efficiently infer parameters when simulations are too costly to be used directly in the inference algorithm. Here we use a similar approach and train Gaussian processes to approximate the distance functions between simulated and observed data (one for each simulation model we consider). We set up the Gaussian processes with Scikit-learn [42] and set the kernel to the sum between a rational quadratic kernel and a white kernel. The process can be summarized as follows:

1. We train Gaussian processes to approximate the distance function between synthetic, observed data and simulated data.
2. This surrogate distance is used by pyABC to evaluate candidate particles. No extra simulations are performed during this process.

Gaussian processes not only estimate the mean value of the distance function at a given point in parameter space, they also estimate the uncertainty of this value. In our case, this is important because of the stochastic aspect of particle acceptance in ABC. Specifically, the same particle may be accepted or rejected depending on the simulated data from this particle, especially if it comes from a stochastic model. Thus, by modeling the stochastic variations around the measured distance with Gaussian processes, we can reproduce this aspect.

Naturally, the GP surrogates come at the cost of an approximation error. Since we do not know the true posterior distribution, it is difficult to quantify what is the effect of this approximation on our results. Järvenpää *et al.* introduced a measure of utility to quantify the goodness of fit of the Gaussian processes. Although in absolute terms, this raw number is not very enlightening, it can be used to compare how the approximation performs in different configurations. In all our experiments, the utility was never correlated with the patterns exhibited on the error maps (see Figure S1), suggesting it had only a low impact on the inference error.

By using this approximate distance, we can greatly reduce the computational cost of running ABC, regardless of the model used to simulate the data. In particular, even detailed models which would be far too computationally expensive to be used as is in ABC can be plugged in into our pipeline. This makes it possible to set a baseline in terms of what could be achieved in terms of accuracy when using simpler models.

## 4 Results

We next use the pipeline described in Section 3 to investigate different scenarios for inferring parameters. We first detail the model used to generate the data in Section 4.1 and then elaborate on how we measure the posterior error in Section 4.2. We then investigate how the amount of data in terms of time sampling density, throughput and observed species (protein, mRNA or both) can influence accuracy and in what situations we clearly benefit from using the higher spatial model fidelity, as motivated by data. We then investigate where in the parameter space each model performs best, and which combination of model and distance metrics gives the best results overall.

### 4.1 GRN models, synthetic data generation and distance metrics

In [11], we studied a negative feedback motif motivated by the Hes1 GRN, in which a gene represses its own expression through a negative feedback loop: mRNA is transcribed in the nucleus and diffuses out into the cytoplasm where it is translated to proteins. These proteins then diffuse back to the nucleus and repress the gene. This process is illustrated in Figure 2. The delay between the moment mRNA is produced and the moment proteins diffuse back into the nucleus and bind to the gene tends to generate oscillations in gene expression level. These chemical reactions are described in Equations 3–7 while their parameters are summarized in Table 1. Three approaches were used to model this network: the first approach consists of a detailed particle model based on the Smoluchowski diffusion limited equations. We use the widely-used software Smoldyn [4] to simulate the model, thus we refer to it simply as *Smoldyn* in the remainder of this paper. The second approach is a cheap multiscale approximation from [11]. Here the cell geometry is divided into two compartments (cytosol and nucleus) which are themselves considered to be well-mixed. Transition rates between these two compartments are then derived using hitting-time analysis on the Smoluchowski model, thus capturing some of the spatial effects. The model is simulated with the standard SSA and is referred to as the Compartment-Based Model (CBM). Finally, the entire cell is considered to be well-mixed and the model (WMM) is simulated using SSA. Note that we use diffusion-limited reaction rates in the association reactions between the gene and protein, thus all three models explicitly involve all physical constants, enabling direct comparison. For all SSA simulations we use the software Gillespy2 in the StochSS suite of tools [30, 13].

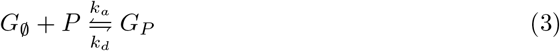

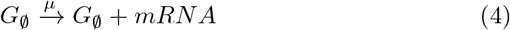

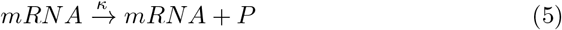

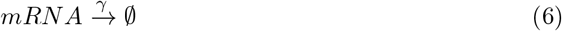

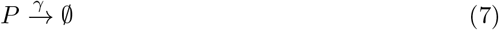

**Figure 2:**
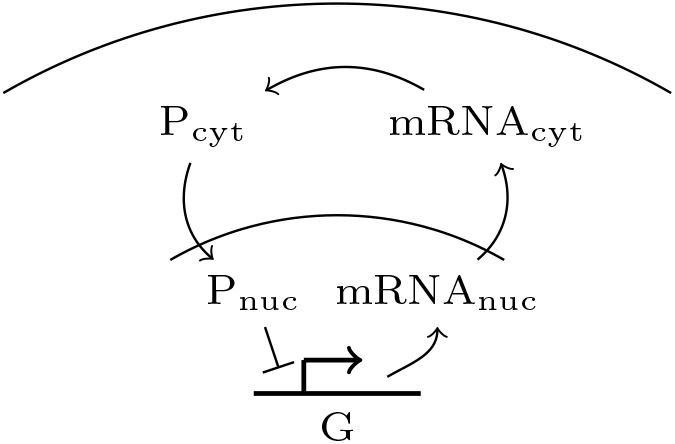
Sketch of the genetic motif studied in this article. A gene, placed at the center of the nucleus of the cell, transcribes mRNA. mRNA then diffuses out of the nucleus and into the cytoplasm, where it is translated to proteins. These proteins then diffuse back into the nucleus, where they repress the expression of the gene.

**Table 1:** Base parameters as presented in [11]. The parameters to be estimated with Bayesian inference are highlighted in grey and are varied over several orders of magnitude in the synthetic data sets. Parameters *μ*, *κ* and *γ* are varied simultaneously by multiplying them by a common variable, noted *χ*. This variable and the diffusion constant are the targets to be inferred.

| Parameter | Description | Localization | Base Value | Unit |
| --- | --- | --- | --- | --- |
| $R$ | cell radius | | 6.0 | $\mu\text{m}$ |
| $r$ | nucleus radius | | 2.5 | $\mu\text{m}$ |
| $D$ | diffusion constant | | 0.6 | $\mu\text{m}^2 \text{min}^{-1}$ |
| $k_a$ | binding rate | nucleus | $1.00 \times 10^9$ | $\text{M}^{-1} \text{min}^{-1}$ |
| $k_d$ | unbinding rate | nucleus | 0.1 | $\text{min}^{-1}$ |
| $\mu$ | transcription rate | nucleus | 3.0 | $\text{min}^{-1}$ |
| $\kappa$ | translation rate | cytoplasm | 1.0 | $\text{min}^{-1}$ |
| $\gamma$ | degradation rate | entire cell | 0.04 | $\text{min}^{-1}$ |

Our goal is to systematically evaluate the quality of parameter estimates inferred with ABC, depending on which of the three models are used to simulate the data, and on the amount of data available. We use the well-established Sequential Monte-Carlo (SMC) variant of ABC as provided in the pyABC Python package [33]. The prior is set to a log-uniform distribution between 0.25 and 16 for *χ* and between 0.0039 and 16 for the diffusion constant. For each setup, we infer parameters from 256 different synthetic data sets generated with Smoldyn, and compare the posterior distributions to the true parameters. This gives us a map of inference performance for the systems ranging from the strongly diffusion-limited regime to the well-mixed regime.

We consider three distance metrics to measure the discrepancy between candidate particles and the observed data:

1. First we consider four common summary statistics, namely the mean value, the minimum value, the maximum value and the standard deviation of a trajectory. We then take the expected value over all trajectories and for both species, and compute the distance between simulated data and synthetic data using the *L*_2_ norm. This setting represents a likely first setting formulated manually by a modeler. We refer to this setting as *naive statistics*.
2. Second, we select optimal summary statistics using the AS algorithm described in Subsection 2.2. The selected statistics are: the longest strike below the mean, the longest strike above the mean, the mean absolute change, the maximum, the minimum and the variance. The distance is computed as in 1 above. We refer to this setting as *optimized statistics*.
3. Third, following [35], we observe the distribution of molecular counts for each species and at every time point, i.e. we compute a histogram density approximation of the cumulative density function (CDF) based on the observed trajectories for each time point, and then compute the average Kolmogorov distance between the two data sets over these distributions. We refer to this setting as *Kolmogorov distance*.

The data used for our study is publicly available as a .json file (as well as all the code used in that study) on GitHub^1^. The data this study is based on is taken from a previous study [11] and is publicly available as a .json file also on GitHub^2^. For each pair of diffusion and reactivity coefficient, the data set contains 64 trajectories over 100 time samples for the two species of interest in the system (namely mRNA and proteins), and for each of three models. A burn in period was used at the beginning of each simulation to make sure all trajectories are uncorrelated from the initial condition.

In the data-scenarios detailed in this study, the pipeline is run separately for each model (WMM, CBM and Smoldyn) and then for each of our 256 synthetic data sets. Figure 3 illustrates this process. In total, 768 inferences are performed for each setup. In each scenario we vary the amount of data and the distance metric used each time. All the computations were run on Rackham, a high performance computing cluster provided by the Multidisciplinary Center for Advanced Computational Science (UPPMAX). Each nodes consists of two 10-core Xeon E5-2630 V4 processors at 2.2 GHz and 128 GB of memory. Running the pipeline for one setup (i.e. 256 3 inferences) took approximately 200 core-hours. Specifically, this is the cost of generating the results as presented e.g., in Figure 4. We emphasize that executing our pipeline is only made possible by the use of Gaussian Processes to approximate the distance function between simulated and observed data.

**Figure 3:**
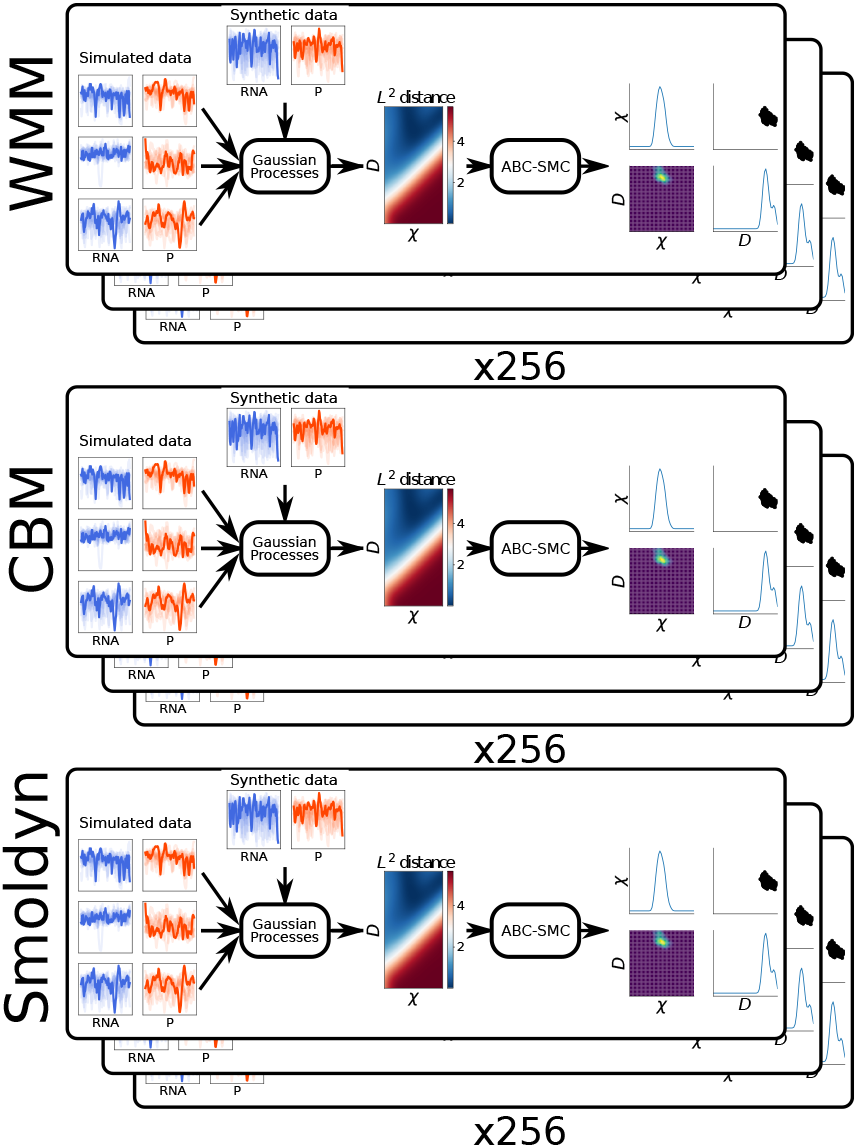
Illustration of the computational experiments performed. For the different data-scenarios and for each combination of distance metric and amount of data, we execute the pipeline using all 256 synthetic data sets as observed data, and for each of the three models.

**Figure 4:**
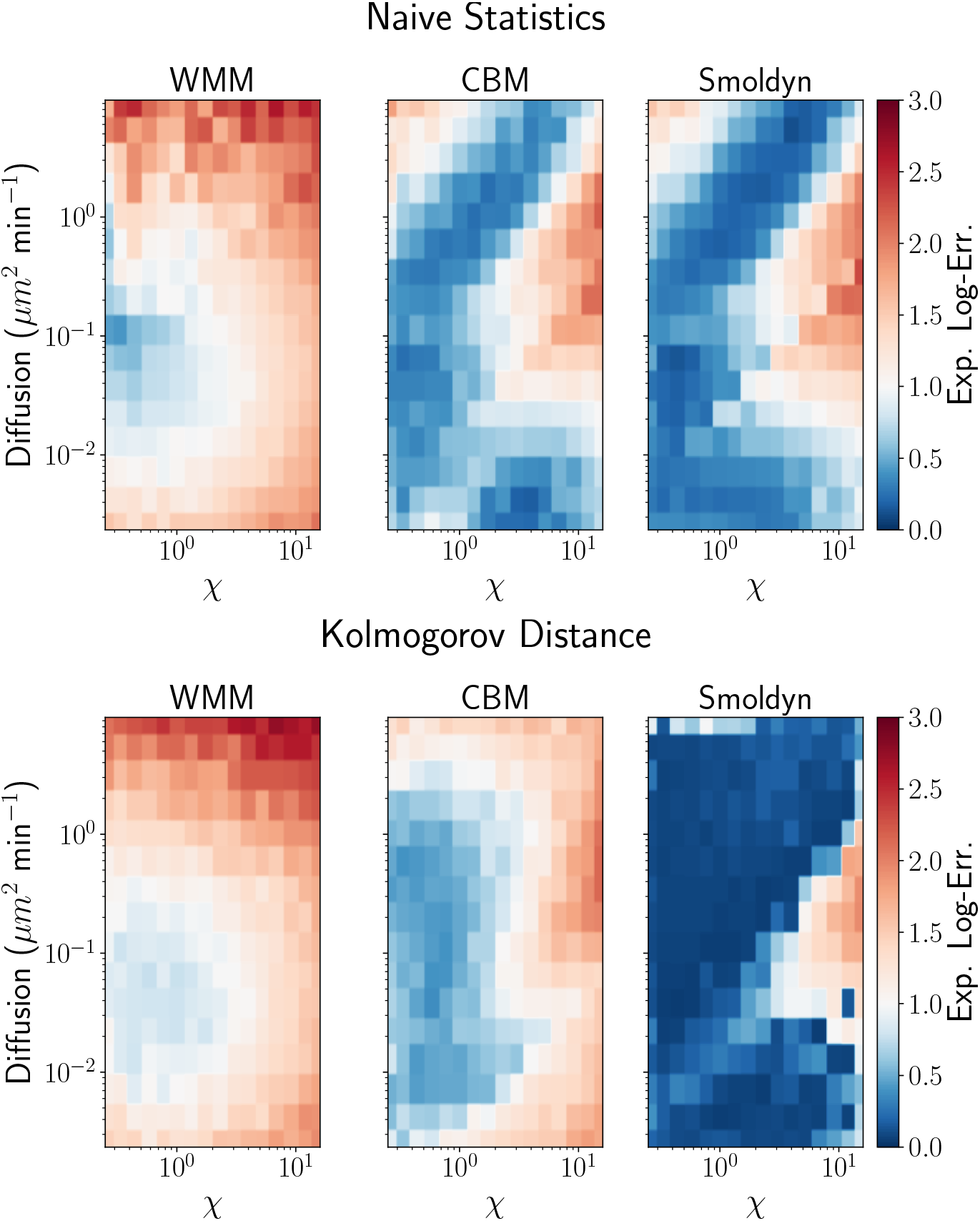
Expected log-error based on summary statistics (upper row) and the Kolmogorov (lower row) distance, for all three models. The CBM performs almost as good as Smoldyn when using summary statistics. When using the Kolmogorov distance, Smoldyn outperforms the two other models.

### 4.2 Depending on distance metric, a detailed spatial model can be motivated also by non-spatial observations

In this section, we compare the performance of the three model levels for parameter inference using the complete observed dataset (64 trajectories sampled at 100 time points) for each of the three distance metrics.

When the true parameters are unknown, it is common practice to report on the posterior mean estimate of the parameters, as well as confidence intervals. If the true parameters are known, as in our case, we can compute the actual error between the parameter estimates and the true parameters, or report if they fall within given confidence intervals. Since the parameter space we use spans several orders of magnitude, computing the relative error in the mean posterior parameters is not very meaningful, i.e., when the true parameters are on the small end of the parameter space, the relative error tends to be large, while if the true parameters are on the bigger end, the relative error will be close to 1. Instead, we choose to report the *expected log-error* with respect to the posterior with the following formula:

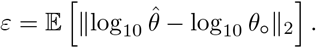

Here *θ*_o_ are the true parameters and 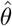 is the estimate of the parameters from retained particles in the posterior, i.e. we compare the estimates in terms of order of magnitudes from the true parameters. This measure also penalizes wide posteriors, which will have a larger expected error than tighter posteriors. This allows us to clearly and concisely report on both the accuracy and uncertainty in the 256 inferences performed in each experiment and for each model. In summary, the chosen error measure takes smaller values for inferences with tight posteriors around the correct expected parameter point and increases both with bias and spread of the posterior (lower inference quality). Fig. 4 shows these error maps for all three models, the naive statistics and the Komogorov distance.

As seen from the figure, we obtain significantly better inference performance when using Smoldyn as the simulator compared to when using the WMM both when using the naive set of statistics and the Kolmogorv distance, even in the well-mixed regions of parameter space. We also see that using the Kolmogorov distance together with the detailed spatial simulator gives the best inference quality overall. This answers one of our initial questions - it is motivated from a inference and data-driven perspective to use models with explicit spatial detail even though observations are more coarse-grained in nature. The nature of observation and the distance metric employed have a large influence here: when using summary statistics rather than Kolmogorov distance, the CBM leads to approximately the same inference quality as Smoldyn, suggesting that the model is able to capture the critical spatial effects on the statistics. We also clearly see the variation of inference quality throughout parameter space – in simple words, some regions are easier to infer than others, and for some regions the expected log error is unacceptably high (2 orders of magnitude or more) even for the best configurations. In particular, with this dataset size we are not able to accurately identify parameters even if using the ground truth model in those regions. We suggest that computing this type of error map for the simulators a modeler considers will also aid when using real experimental data – if the inferred parameter falls in a region of the map where the error is large it is a good indication that care needs to be taken in interpreting the results.

Although the error map in Fig 4 provides a detailed view of the regions where the inference has the lowest error, the level of detail makes it hard to compare two models quantitatively. For instance, in Figure 4, it is unclear how much more accurate Smoldyn is compared to the CBM. Thus, when comparing results for two different settings, we build an enhanced box plot. Contrary to the regular box plot, where only the quartiles are shown, the extended box plot also displays the next 2*^n^*-quantiles above the upper quartile and below the lower quartile, thus giving a better view on the distribution of tail values [26]. Figure 5 illustrates such visualization technique. By representing the error in this way, we trade local information in parameter space for easier, global comparison.

**Figure 5:**
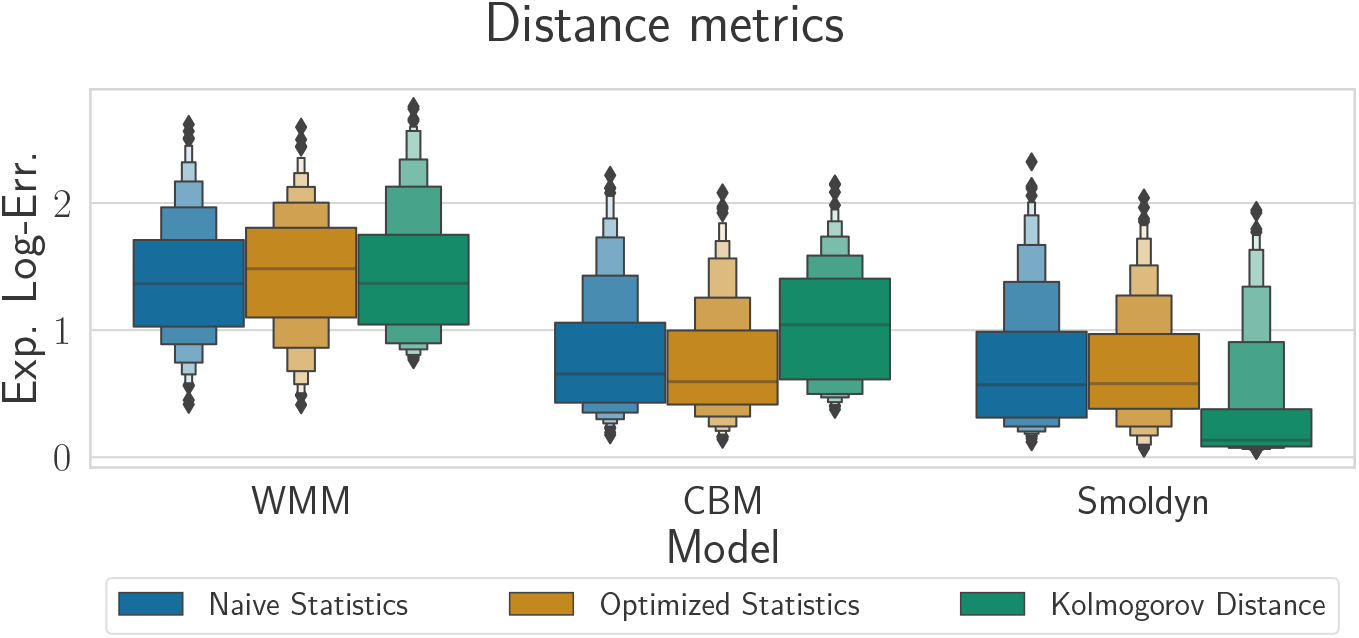
Expected log-error for all three models and all three distance metrics. There is general convergence when adding more details to the model, although for a given model, no metric is consistently more accurate than the others.

Here we also show results using the optimized statistics. Although they do improve inference for the synthetic data sets where it was already accurate with the naive summary statistics (Figure S3), overall, they do not bring significant improvements in inference quality, and will in some cases also lead to worse performance than the naive set. This illustrates the challenge in choosing good statistics.

For the WMM, inference quality does not depend on the distance metric used, suggesting that model error is dominating. When it comes to the Kolmogorov distance, we see minor improvements when going from the WMM to the CBM, and indeed distance metrics based on summary statistics outperform the Kolmogorov distance for the CBM. This can be understood in light of the approximation properties of the CBM: in [11], we showed that, when using summary statistics, the CBM could better approximate Smoldyn than when using Kolmogorov distance. There is a significant improvement in using the CBM versus the WMM. When it comes to Smoldyn, however, the Kolmogorov distance largely outperforms both naive and optimized summary statistics. In fact, this is the only case where the parameters were inferred with acceptable accuracy over the majority of the parameter space. This suggests that this distance is more robust to outliers than summary statistics based metrics. Thus, when combined with an accurate model, this distance is capable of producing more accurate results when inferring the parameters.

We conclude with a comment on the accuracy of inference versus the local approximation quality of the coarse grained model alternatives. In [11], we showed that, when using summary statistics, the CBM could accurately approximate Smoldyn throughout the entire parameter space considered in this study. When using the Kolmogorov distance, we showed that it was only highly accurate in the upper left half of the parameter space, namely when diffusion is high and chemical reactions are slow. Regardless which distance metric was used, the WMM was only accurate in the upper left half of the parameter space. Inspecting Figure 4 and comparing it to the Figure 3 in [11], we can see that the accuracy of inference does not directly depend on the accuracy of the coarse-grained model at location of the true parameters, i.e. inference can be relatively accurate when the coarse-grained model is not a good approximation and it can also be very inaccurate even when the coarse-grained model *is* a good approximation. That being said, we can still see that parameter inference becomes more accurate when a more detailed model is used, regardless of the distance metric in use. All in all, this shows that accurate inference does not depend only on how good the coarse-grained model fits the detailed spatial model at the true parameters, but rather how well it globally fits the fully detailed model over the parameter space.

### 4.3 FACS-like data with Kolmogorov distance is able to discriminate between low-and high model fidelities even for coarse time samples

In this section, we compare the performance of our three models in terms of parameter inference in a fluorescence activated cell sorting (FACS) like setting. First developed in the 1960s, flow cytometry is a popular analytical cell-biology technique that utilises light to count and profile cells in a heterogenous fluid mixture. Flow cytometry is a particularly powerful tool because it allows a researcher to rapidly and accurately acquire population data related to many parameters from a heterogeneous fluid mixture containing live cells. Flow cytometry is used extensively throughout the life and biomedical sciences, and can be applied in any scenario where a researcher needs to rapidly profile a large population of loose cells in a liquid media. FACS differs from conventional flow cytometry in that it allows for the physical separation, and subsequent collection, of single cells or cell populations [32]. FACS is useful for applications such as establishing cell lines carrying a transgene, enriching for cells in a specific cell cycle phase, or studying the transcriptome, or genome, or proteome, of a whole population on a single cell level.

In order to mimic a FACS experimental setup, we followed the method of [35]. In [35] measurements are taken at regular intervals. For a given interval, the Kolomogorov distance between the observed distribution and the simulated distribution is computed and the average distance across all measurements is reported. Our data set contains a total of 100 time measurements (one every ten minutes). We reduce this data set to contain only 12, 6 and 3 measurements. We then run our inference pipeline on each coarsened data set with each model. The expected log-error is reported in Figure 6.

**Figure 6:**
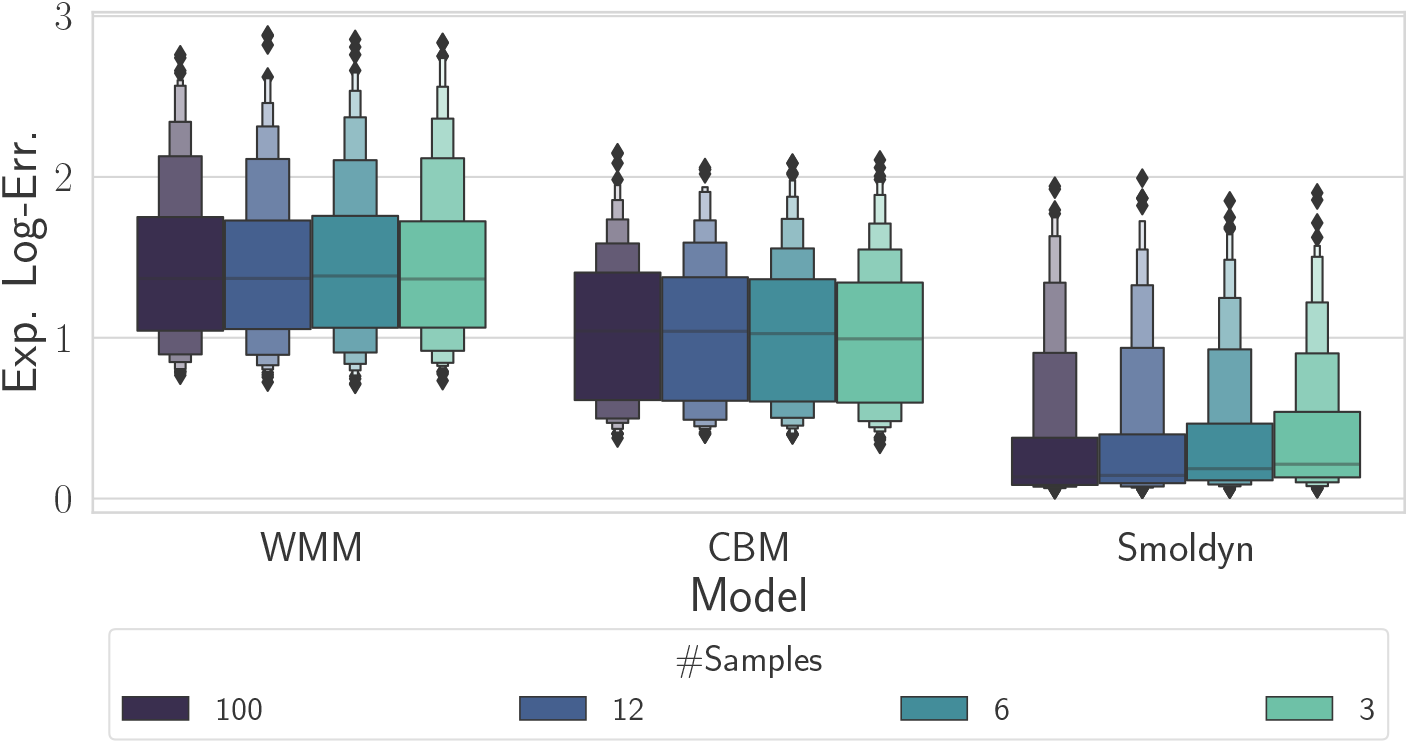
Expected log-error based on the Kolmogorov distance when increasing the number of time samples from 3 to 100. The error only decreases significantly when Smoldyn is used, suggesting model error is the limiting factor in the case of the WMM and the CBM.

Strikingly, we found that, while Smoldyn had the lowest error, reducing the amount of data available had only little effect when using the WMM or the CBM, suggesting that model error was the limiting factor. Although the CBM achieved a lower error level than the WMM, it did not improve with larger numbers of samples, suggesting again that the model is not able to accurately capture the data. Put another way, there was no statistically significant improvement in the error distribution across the considered parameter space for 3 samples versus 100 samples. Increasing the number of time samples only had some effect in the case of Smoldyn. Overall the results of this section suggest that using more time samples is only beneficial if enough computational power is available to use the detailed model.

### 4.4 Protein measurements are more important for inference accuracy than mRNA measurements

Depending on time and/or budgetary constraints, researchers may have access to mRNA data, protein data or both mRNA and protein data. While some dynamical models have been used to infer networks using only mRNA data [39], others have been constrained using both mRNA and protein data [46]. However, to the best of our knowledge the relative importance of mRNA and/or protein data for model inference is not well studied.

In this section, we compare the performance of our three models in three different data scenarios using three different distance measures. In terms of data, we compare using only mRNA data, using only protein data or using both mRNA and protein data. In terms of distance measures, we compare naive statistics, optimized statistics and the Kolmogorov distance metric. We present our findings for this section in Figure 7.

**Figure 7:**
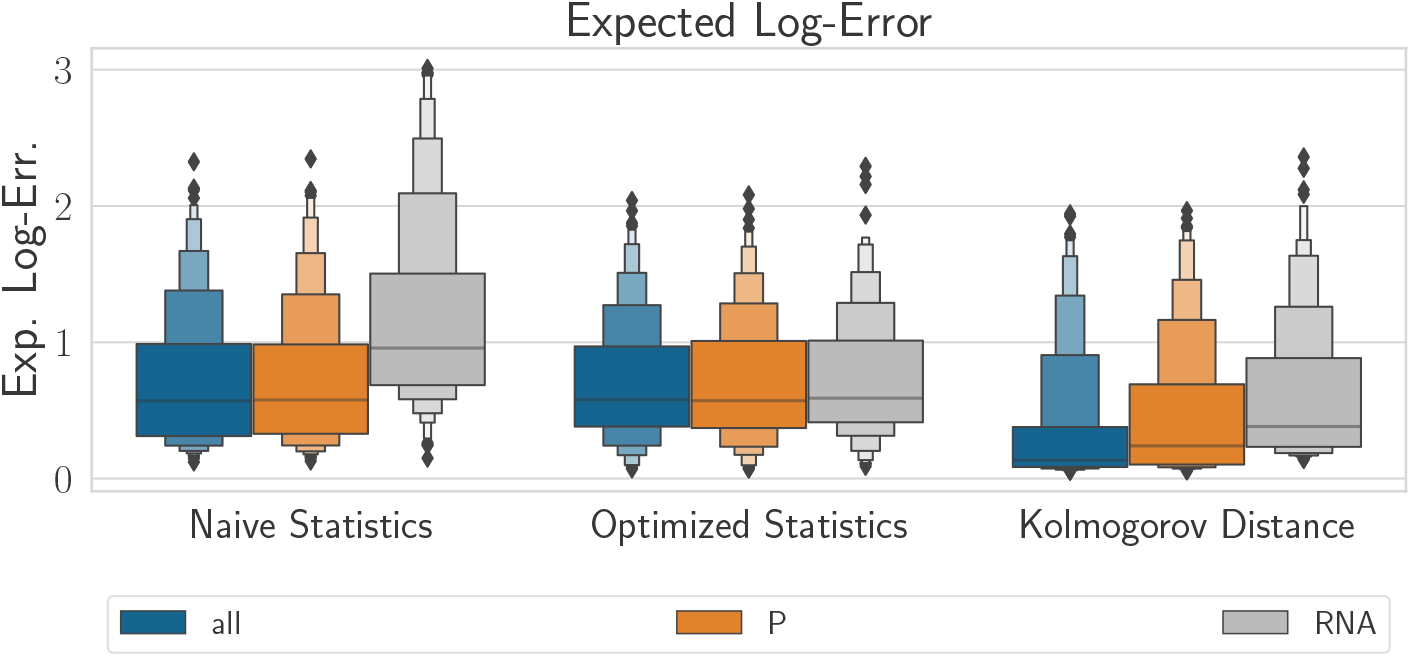
Expected log-error when using all species, only proteins or only mRNA based on all three distance metrics and Smoldyn. Using only RNA tends to decreases the accuracy of the inference. Figure S2 shows the same plots with the CBM and the WMM.

Analyzing data simulated using Smoldyn, we found that only measuring mRNA levels had worse accuracy than measuring only protein levels or measuring both mRNA and protein levels. We also found that collection of mRNA and protein data was only beneficial when used in conjunction with the Kolmogorov distance. In contrast, we found that data simulated using the WMM or CBM showed little difference between all three distance metrics for all three data scenarios (Figure S2). The results here suggest that model error is the limiting factor with respect to inference accuracy and that only measuring one species is enough if only the WMM or CBM models can be used.

## 5 Discussion

In this manuscript, we built a computational pipeline to systematically investigate the performance of ABC with respect to the choice of model, the nature and amount of observed data and the way we compare true and simulated data in likelihood-free inference with ABC (i.e., the distance metric). We applied this pipeline to a canonical model of negative feedback regulation and several experimentally motivated data-scenarios. We showed that the pipeline can be used to reveal insights into which combination of model, amount of data and distance metric can be expected to lead to the most accurate inference result. When analyzing inference error over the parameter space, one can then identify areas where inference is most likely to be accurate when executed against experimental data. Using the complete set of observations (64 trajectories each with 100 equidistant temporal observations) we found that parameter inference was most accurate overall using the most detailed model (as expected) and the Kolmogorov distance (as opposed to summary statistics).

One key question we sought to answer using our simulation-driven inference pipeline was under which inference conditions we were able to see a clear benefit from using the full spatial model (Smoldyn). Since the true synthetic data was generated with Smoldyn we expected that, if inference conditions allowed, using that simulator should lead to superior inference accuracy. In our experiments, we observed a clear benefit from using Smoldyn over the WMM model when using both naive and optimized summary statistics (Fig 5). However, when using these summary statistics, there was no significant benefit in using Smoldyn over a simpler multiscale compartment model that incorporates some spatial features of the simulation. However, when distribution data was used with the Kolmogorov distance metric instead of the mean values, only the fully detailed spatial model managed to leverage the information in the data (Fig. 5 and Fig S4).

From an experimental point of view, another interesting question we sought to answer using our pipeline is whether it is better for inference accuracy to observe mRNA, protein, or both. We ran experiments for this scenario and showed that, for the negative feedback model we consider, only measuring mRNA levels gave the least accurate results. Measuring only proteins was usually as accurate as measuring both species except when Smoldyn and the Kolmogorov distance were used, in which case a relatively small gain was obtained from observing both species.

We considered two sets of summary statistics, a naive pool consisting of typical statistics a modeler might choose for a first attempt at inference, and a set of statistics obtained by state-of-the art summary statistic selection from a larger pool of time series features. These optimized summary statistics did not show major improvements compared to the naively chosen statistics, highlighting the fact that statistics selection is a challenging problem. It is possible, of course, that there exists a possible set of statistics that performs better. However, in our experiments the Kolmogorov distance approach gave more robust results. Here, using Kolmogorov distance to compare summary statistics (as opposed to taking expectations) shows a gain for the spatial model, however the standard Kolmogorov distance measure led to more accurate results still, see Fig. S4. An interesting future direction would be to also compare to an emerging class of methods that automatically learn good or optimal statistics by training a regression model [17] to predict the posterior mean parameters [2, 29, 58]. This approach is particularly useful in cases where optimal summary statistics may not even exist in the candidate pool to select from, but comes with an overhead of requiring a sizeable amount of training data (pairs of the form - **y**, *θ*) to obtain the regression models.

Executing the pipeline over such a large section of the parameter space we used was only made computationally tractable by the use of Gaussian processes to approximate the distance metric when assessing the discrepancy of candidate particles. Indeed, as shown in [11], inferring the parameters for a single synthetic data set with the WMM or the CBM took between 10 and 100 core-hours, and a lower bound estimate for running ABC using Smoldyn ranged from 780 to 4635 core-hours depending on parameter values. Clearly, running this process on a large amount of synthetic data sets and in various configurations in terms of distance metrics and amount of data is not computationally tractable. In comparison, inferring the parameters of one data set with the Gaussian process approximation took less than one core-hour including the time to train the Gaussian processes. There is of course some error associated with using such an approximation, and this error should be monitored to differentiate it from other sources. In Figure S1, we looked at the utility as defined by Järvenpää *et al.* [27] and showed that it was relatively constant, suggesting our results are consistent over the parameter space. In a further study, we will use this pipeline to estimate the error stemming from the Gaussian processes by varying the amount of training data and studying the effect it has on the posterior distribution. Executing this pipeline should only be seen as a preliminary step to select the components involved in parameter inference. Once the setup has been calibrated and experimental data has been collected, regular ABC can be used, provided simulating the model is not too computationally expensive.

In this study, we used the model of highest fidelity when it comes to biophysical realism as a proxy for the ground truth. This enabled us to compare different model levels to each other. When attempting to make extrapolations of the pipeline results to real experimental conditions, it is important to recall that in practice, real experimental data will be noisier than simulated data, first because every model is wrong (although some of them are useful, as the quote says), and then because even if the model is an accurate representation of reality there will always be measurement noise due to limitations in the experimental protocol. Here we opted for “perfect synthetic data” to make interpretation of numerical settings and model comparison easier, however if would be possible and interesting to repeat the numerical experiments using a measurement error model. In this way it would be possible to also study the different settings and scenarios (e.g. summary statistics vs. Kolmogorov distance) with respect to robustness to noise.

In an inference scenario using real experimental data the “true” parameters are unknown and it will be difficult to validate parameter inference, given that the model error is unknown. As discussed in the introduction, most often a modeler will favor one model type. As our study revealed however, it can be important to compare different models to each other and in particular compare different inference strategies.

Bayesian parameter inference is an efficient technique to find areas of the parameter space that resembles observed data. If the parameters are identifiable, the estimated parameters will correspond to the “true” parameters. Showing identifiability, however, is only possible for a limited class of models [47]. Our approach provides an alternative where it is possible to quantify potential error due to model choice, summary statistics or lack of data in the context of Bayesian inference.

Finally, we note that in this work we were interested in using the model to accurately learn the underlying model parameters. For a given set of observed data and for given numerical inference settings, it is then clear that a modeler should favor the computationally cheapest model that results in good parameter inference. Note here that, due to practical aspects of likelihood-free inference it is not enough if a coarse-grained approximation is accurate only at the true parameter point, it needs to be accurate throughout the support of the prior distribution. Given that true parameters are unknown in practice, this means that we either need some prior knowledge about in which regimes the true parameter will fall (in which case we can use a pipeline like ours to suggest the stability of inference to different model choices), or we need to seek a globally accurate approximation. We emphasize that this problem is different from a typical ABC-based Bayesian model selection problem, in which we seek to use simulators for different models to compute the probability of the models generating the observed data. In that setting we allow models to take “wrong” physical parameter values as long as that model configuration is capable of generating trajectories close to the observed data. To illustrate this, we used pyABC to solve a model selection problem comparing the WMM, CBM and Smoldyn for each of the synthetic datasets using either the naive summary statistics or the Kolmogorov distance (Fig. 8). As can be seen, when using summary statistics we obtain an intuitive picture where Smoldyn is clearly favored in the highly diffusion limited regime, the CBM is favored in a transition region, and all models are about equally probable as the systems becomes more well-mixed (Fig. 8 upper row). However, using the Kolmogorov distance, which we showed gave superior inference accuracy, the WMM model is favored almost everywhere in the parameter space (Fig. 8 lower row) even though the Smoldyn simulations leads to lower inference error for most parameter combinations, c.f. Fig. 5. This highlights that care needs to be taken both when interpreting model selection results, not to mix up objectives (inferring true parameters vs. a model that can generate the observed data) and when choosing ABC setup since a setup that is robust for accurate inference can be non-robust for Bayesian model selection.

**Figure 8:**
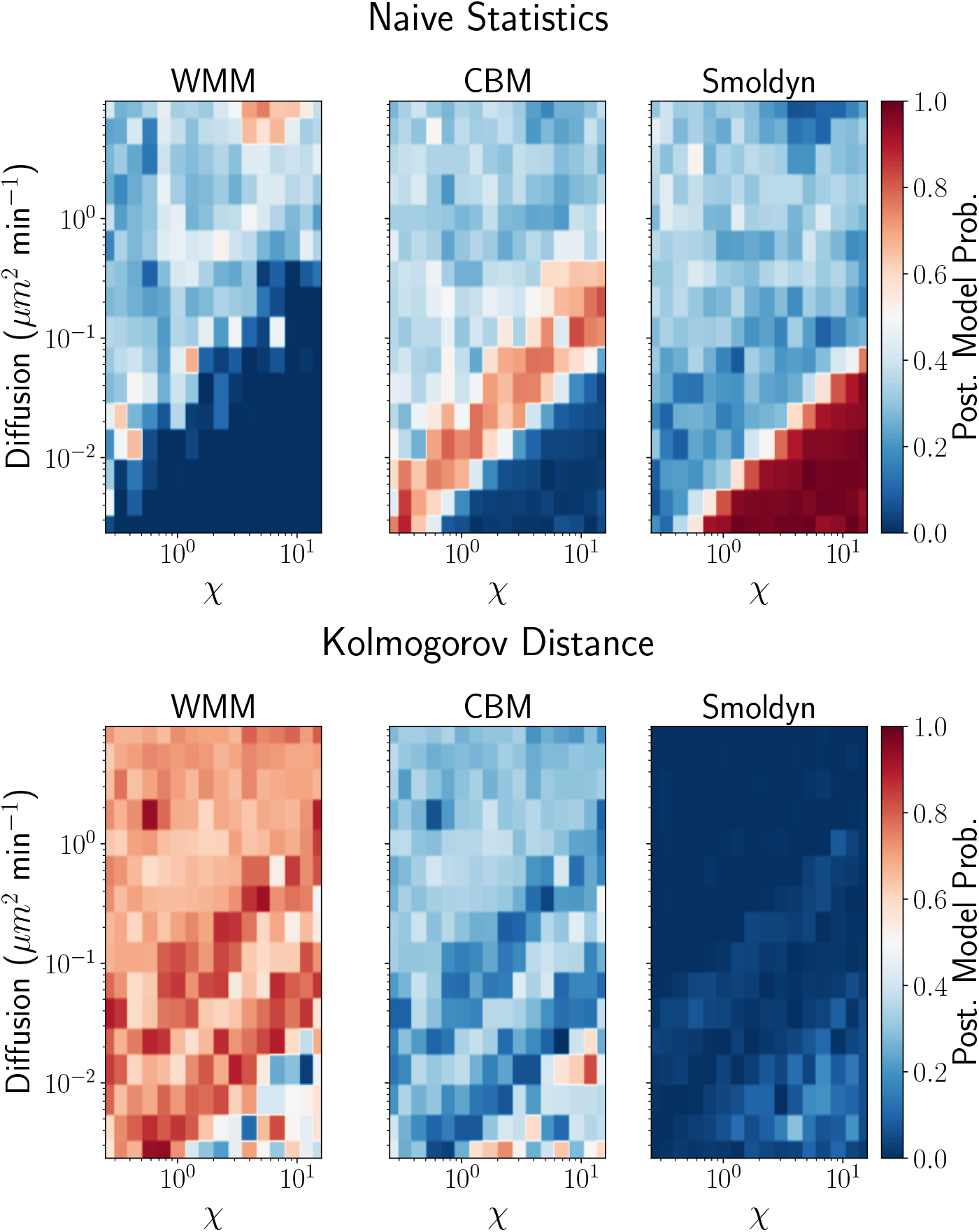
Model probabilities as computed by pyABC when comparing the three models. The algorithm adds the model index as a new parameter to be inferred, the model probabilities are then the posterior distribution over this parameter.

## 6 Acknowledgments

This work has been funded by the Swedish research council (VR) under award no. 2015-03964, by the eSSENCE strategic collaboration of eScience, and by the NIH under grant no. NIH/2R01EB014877-04A1. The computations were performed on resources provided by SNIC through Uppsala Multidisciplinary Center for Advanced Computational Science (UPPMAX) under Project 2019/8-227.

## 7 Supplementary Material

**Figure S1:**
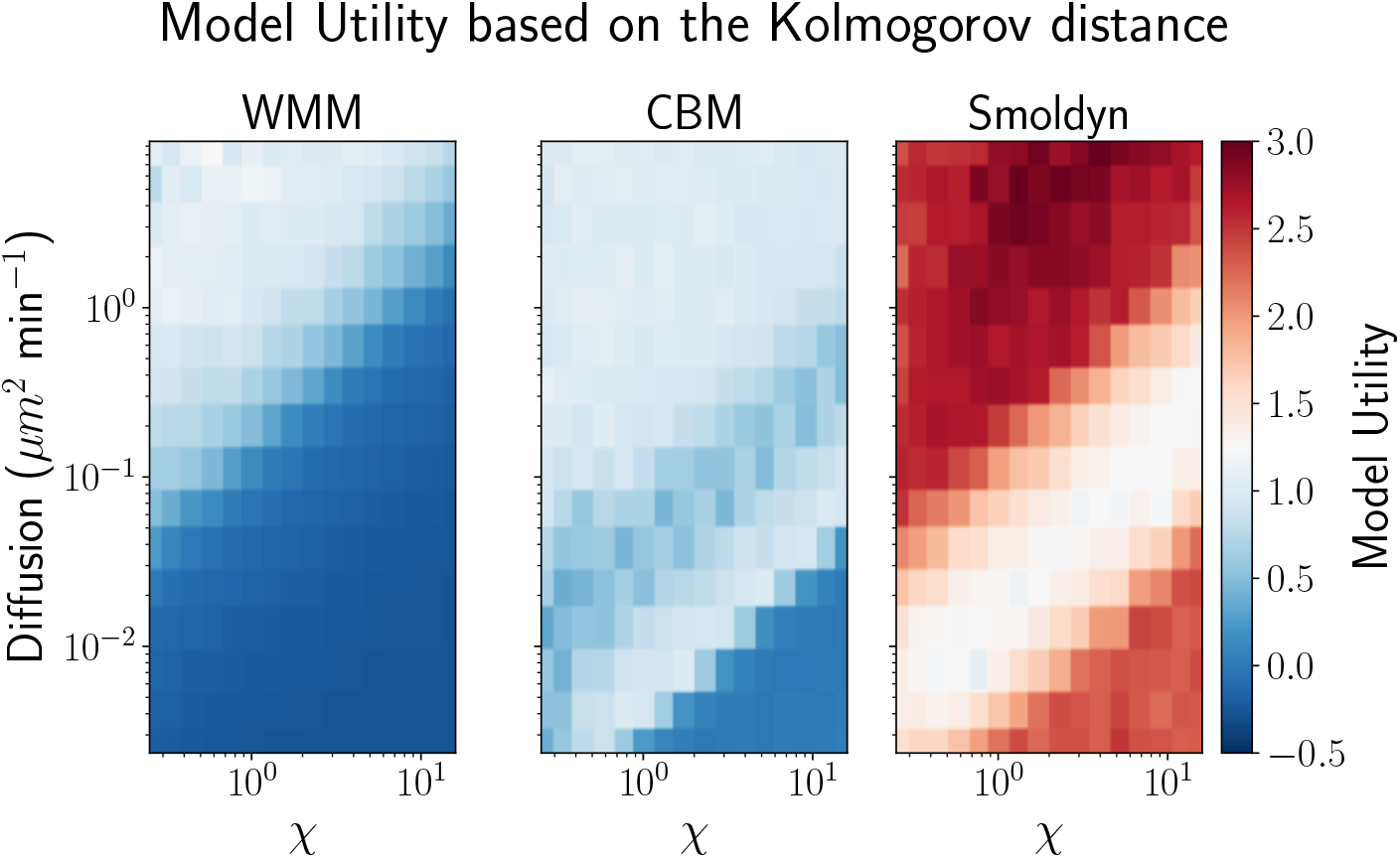
Utility map for all three models when using the Kolmogorov distance. Overall, it does not correlate with the error estimates presented in Figure 4.

**Figure S2:**
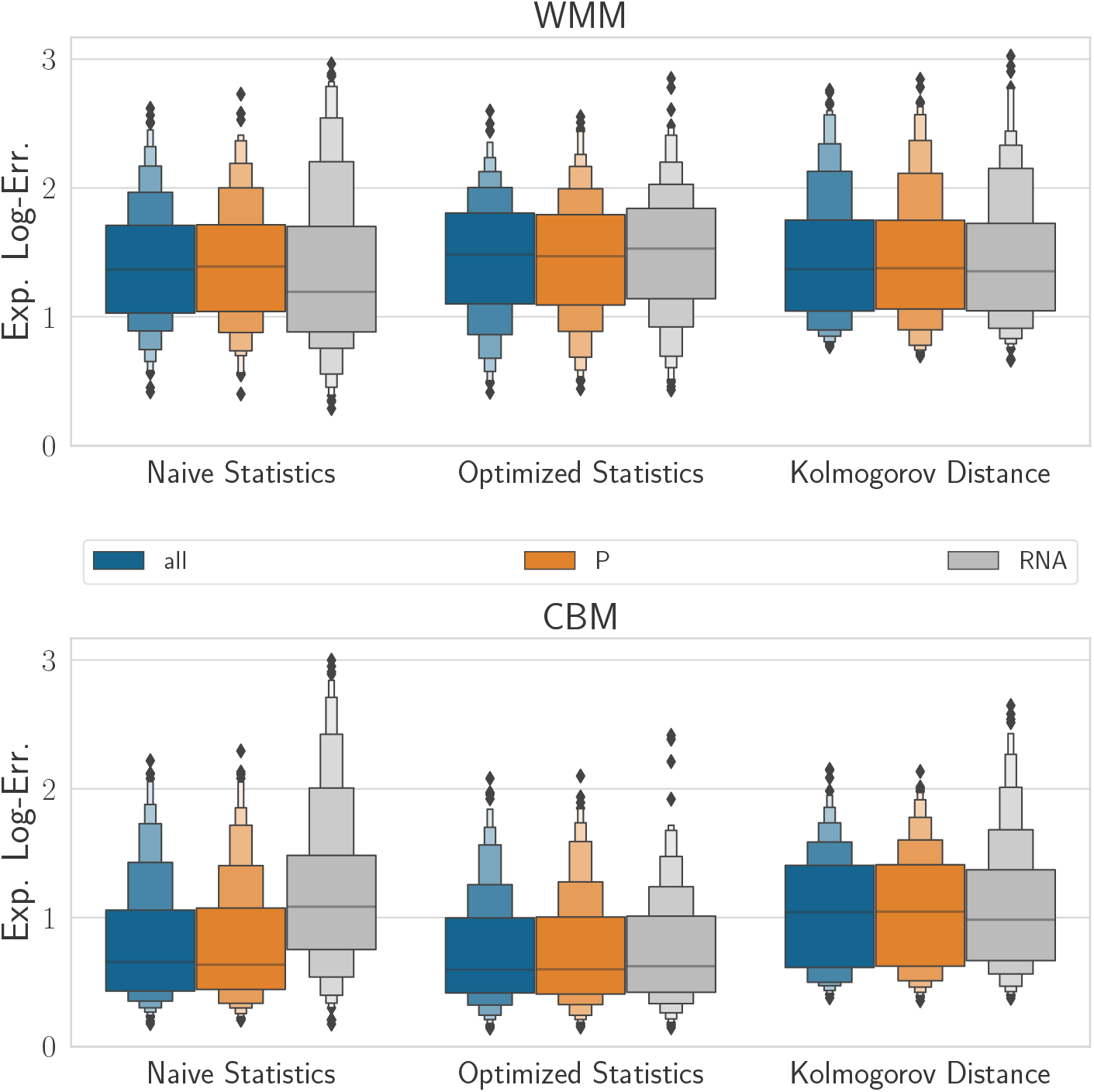
Comparison of expected log-error for the WMM and the CBM, for all three different distance metrics and when measuring only protein levels, only mRNA levels, or both. Overall no big difference can be observed, contrary to the case where the Smoldyn was used.

**Figure S3:**
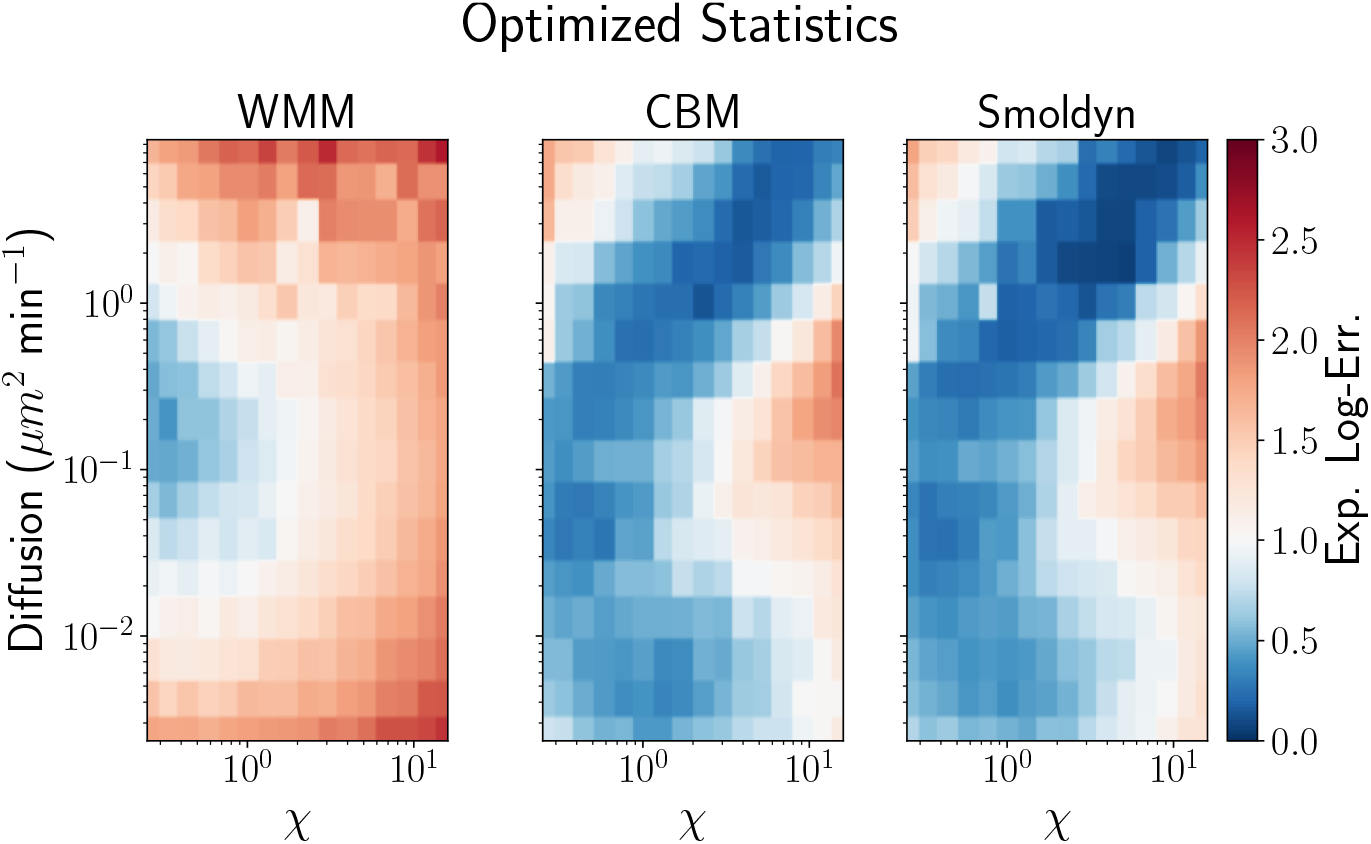
Expected log-error for all three models based on optimized summary statistics. The error is slightly lower than when only basic summary statistics are used.

**Figure S4:**
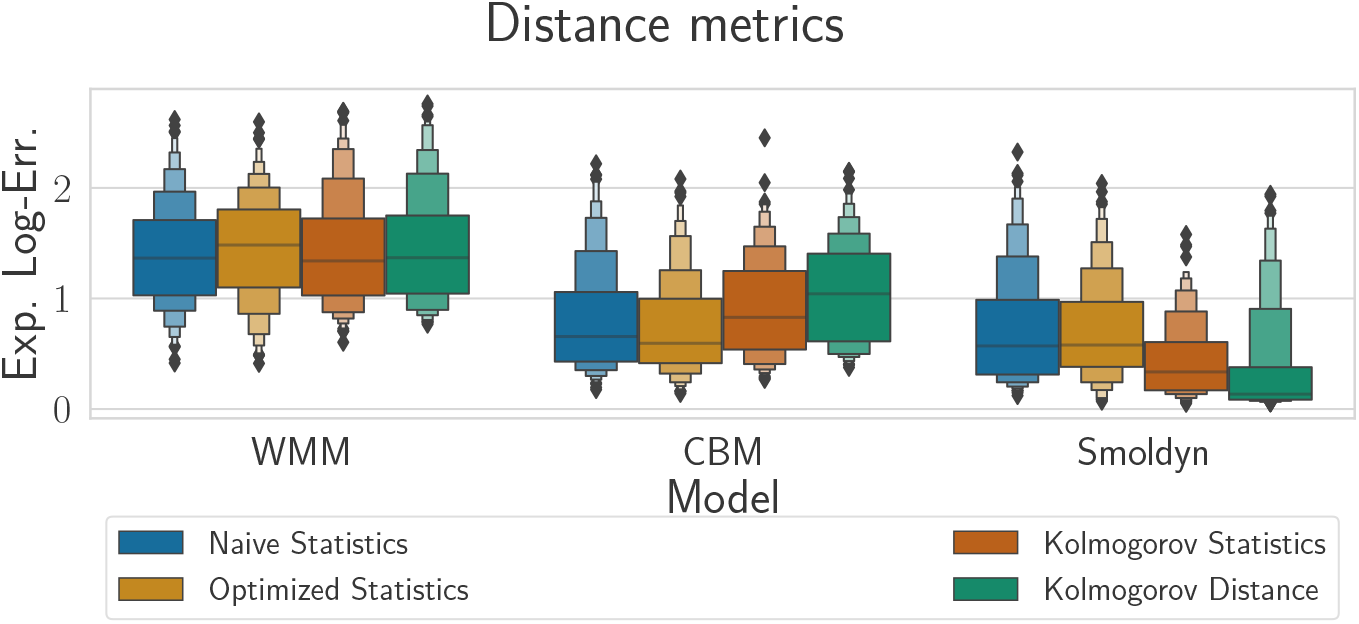
Comparison of distance measure based on expected value or based on distribution. When comparing simulated and true data via summary statistics for datasets with multiple trajectories the most straightforward and the most common way is to compute the statistic for each trajectory and then compare the expected values for true and simulated data. This is what is done in the main manucript when using summary statistics. As a more elaborate alternative we can also compare the summary statistics using the Kolmogorov distance (Kolmogorov statistics). This entails computing the histogram CDF for the statistic at each timepoint, then taking the Kolmogorov distance between true and observed data. As can be seen, this approach is advantageous when using the Smoldyn simulator, however it does not lead to as low errors as directly comparing the copy numbers as done in the main manuscript (Kolmogorov distance). For the CBM model, however, taking a distribution measure leads to higher error.

## Footnotes

1 https://github.com/Aratz/PipelineforParameterInference

2 https://github.com/Aratz/MultiscaleCompartmentBasedModel/blob/master/data/data.zip

## Notes

https://github.com/Aratz/PipelineforParameterInference

### Competing Interest Statement

The authors have declared no competing interest.

### Summary of Updates

We have fixed some typographical errors and made the text clearer in some parts of the manuscript.

## References

[1] Ruedi Aebersold and Matthias Mann. Mass spectrometry-based proteomics. Nature, 422(6928):198–207, 2003.

[2] Mattias Åkesson, Prashant Singh, Fredrik Wrede, and Andreas Hellander. Convolutional neural networks as summary statistics for approximate bayesian computation. arXiv preprint arXiv:2001.11760, 2020.

[3] Cem Albayrak, Christian A Jordi, Christoph Zechner, Jing Lin, Colette A Bichsel, Mustafa Khammash, and Savaş Tay. Digital quantification of proteins and mrna in single mammalian cells. Molecular cell, 61(6):914–924, 2016.

[4] Steven S Andrews, Nathan J Addy, Roger Brent, and Adam P Arkin. Detailed simulations of cell biology with smoldyn 2.1. PLoS Comput Biol, 6(3):e1000705, 2010.

[5] Sean C Bendall, Erin F Simonds, Peng Qiu, D Amir El-ad, Peter O Krutzik, Rachel Finck, Robert V Bruggner, Rachel Melamed, Angelica Trejo, Olga I Ornatsky, et al. Single-cell mass cytometry of differential immune and drug responses across a human hematopoietic continuum. Science, 332(6030):687–696, 2011.

[6] Samuel Bernard, Branka Ĉajavec, Laurent Pujo-Menjouet, Michael C Mackey, and Hanspeter Herzel. Modelling transcriptional feedback loops: the role of gro/tle1 in hes1 oscillations. Philosophical Transactions of the Royal Society A: Mathematical, Physical and Engineering Sciences, 364(1842):1155–1170, 2006.

[7] Alexander P Browning, David J Warne, Kevin Burrage, Ruth E Baker, and Matthew J Simpson. Identifiability analysis for stochastic differential equation models in systems biology. Journal of the Royal Society Interface, 17(173):20200652, 2020.

[8] Bogdan Budnik, Ezra Levy, Guillaume Harmange, and Nikolai Slavov. Scope-ms: mass spectrometry of single mammalian cells quantifies proteome heterogeneity during cell differentiation. Genome biology, 19(1):1–12, 2018.

[9] Kevin Burrage, Pamela M Burrage, André Leier, Tatiana Marquez-Lago, and Dan V Nicolau. Stochastic simulation for spatial modelling of dynamic processes in a living cell. In Design and Analysis of Biomolecular Circuits, pages 43–62. Springer, 2011.

[10] Mark Chaplain, Mariya Ptashnyk, and Marc Sturrock. Hopf bifurcation in a gene regulatory network model: Molecular movement causes oscillations. Mathematical Models and Methods in Applied Sciences, 25(06):1179–1215, 2015.

[11] Adrien Coulier, Stefan Hellander, and Andreas Hellander. A multiscale compartment-based model of stochastic gene regulatory networks using hitting-time analysis. The Journal of Chemical Physics, 154(18):184105, 2021.

[12] Spyros Darmanis, Caroline Julie Gallant, Voichita Dana Marinescu, Mia Niklasson, Anna Segerman, Georgios Flamourakis, Simon Fredriksson, Erika Assarsson, Martin Lundberg, Sven Nelander, et al. Simultaneous multiplexed measurement of rna and proteins in single cells. Cell reports, 14(2):380–389, 2016.

[13] Brian Drawert, Andreas Hellander, Ben Bales, Debjani Banerjee, Giovanni Bellesia, Bernie J Daigle Jr, Geoffrey Douglas, Mengyuan Gu, Anand Gupta, Stefan Hellander, et al. Stochastic simulation service: bridging the gap between the computational expert and the biologist. PLoS computational biology, 12(12):e1005220, 2016.

[14] Johan Elf, Andreas Doncic, and Mans Ehrenberg. Mesoscopic reaction-diffusion in intracellular signaling. In Fluctuations and noise in biological, biophysical, and biomedical systems, volume 5110, pages 114–124. International Society for Optics and Photonics, 2003.

[15] Ján Eliaš, Luna Dimitrio, Jean Clairambault, and Roberto Natalini. The dynamics of p53 in single cells: physiologically based ode and reaction–diffusion pde models. Physical biology, 11(4):045001, 2014.

[16] Heiko Enderling and Mark AJ Chaplain. Mathematical modeling of tumor growth and treatment. Current pharmaceutical design, 20(30):4934–4940, 2014.

[17] Paul Fearnhead and Dennis Prangle. Constructing summary statistics for approximate bayesian computation: semi-automatic approximate bayesian computation. Journal of the Royal Statistical Society: Series B (Statistical Methodology), 74(3):419–474, 2012.

[18] Zachary R Fox and Brian Munsky. The finite state projection based fisher information matrix approach to estimate information and optimize single-cell experiments. PLoS computational biology, 15(1):e1006365, 2019.

[19] Daniel T Gillespie. A general method for numerically simulating the stochastic time evolution of coupled chemical reactions. Journal of computational physics, 22(4):403–434, 1976.

[20] Daniel T Gillespie. Exact stochastic simulation of coupled chemical reactions. The journal of physical chemistry, 81(25):2340–2361, 1977.

[21] Daniel T Gillespie, Andreas Hellander, and Linda R Petzold. Perspective: Stochastic algorithms for chemical kinetics. The Journal of chemical physics, 138(17):05B201 1, 2013.

[22] Ryan N Gutenkunst, Joshua J Waterfall, Fergal P Casey, Kevin S Brown, Christopher R Myers, and James P Sethna. Universally sloppy parameter sensitivities in systems biology models. PLoS Comput Biol, 3(10):e189, 2007.

[23] Jonathan U Harrison and Ruth E Baker. The impact of temporal sampling resolution on parameter inference for biological transport models. PLoS computational biology, 14(6):e1006235, 2018.

[24] Elizabeth A. Heron, Bärbel Finkenstädt, and David A. Rand. Bayesian inference for dynamic transcriptional regulation; the Hes1 system as a case study. Bioinformatics, 23(19):2596–2603, 07 2007.

[25] Hiromi Hirata, Shigeki Yoshiura, Toshiyuki Ohtsuka, Yasumasa Bessho, Takahiro Harada, Kenichi Yoshikawa, and Ryoichiro Kageyama. Oscillatory expression of the bhlh factor hes1 regulated by a negative feedback loop. Science, 298(5594):840–843, 2002.

[26] Heike Hofmann, Karen Kafadar, and Hadley Wickham. Letter-value plots: Boxplots for large data. Technical report, had.co.nz, 2011.

[27] Marko Järvenpää, Michael U Gutmann, Aki Vehtari, Pekka Marttinen, et al. Gaussian process modelling in approximate bayesian computation to estimate horizontal gene transfer in bacteria. Annals of Applied Statistics, 12(4):2228–2251, 2018.

[28] MH Jensen, Kim Sneppen, and G Tiana. Sustained oscillations and time delays in gene expression of protein hes1. Febs Letters, 541(1–3):176–177, 2003.

[29] Bai Jiang, Tung-Yu Wu, Charles Zheng, and Wing H. Wong. Learning summary statistic for approximate bayesian computation via deep neural network. Statistica Sinica, 27(4):1595–1618, 2017.

[30] Richard Jiang, Bruno Jacob, Matthew Geiger, Sean Matthew, Bryan Rumsey, Prashant Singh, Fredrik Wrede, Tau-Mu Yi, Brian Drawert, Andreas Hellander, et al. Epidemiological modeling in stochss live! Bioinformatics, 2021.

[31] Paul Joyce and Paul Marjoram. Approximately sufficient statistics and bayesian computation. Statistical applications in genetics and molecular biology, 7(1), 2008.

[32] MH Julius, T Masuda, and LA Herzenberg. Demonstration that antigen-binding cells are precursors of antibody-producing cells after purification with a fluorescence-activated cell sorter. Proceedings of the National Academy of Sciences, 69(7):1934–1938, 1972.

[33] Emmanuel Klinger, Dennis Rickert, and Jan Hasenauer. pyabc: distributed, likelihood-free inference. Bioinformatics, 34(20):3591–3593, 2018.

[34] Jochen Kursawe, Ruth E Baker, and Alexander G Fletcher. Approximate bayesian computation reveals the importance of repeated measurements for parameterising cell-based models of growing tissues. Journal of theoretical biology, 443:66–81, 2018.

[35] Gabriele Lillacci and Mustafa Khammash. The signal within the noise: efficient inference of stochastic gene regulation models using fluorescence histograms and stochastic simulations. Bioinformatics, 29(18):2311–2319, 2013.

[36] Jing Lin, Christian Jordi, Minjun Son, Hoang Van Phan, Nir Drayman, Mustafa Fatih Abasiyanik, Luke Vistain, Hsiung-Lin Tu, and Savaş Tay. Ultra-sensitive digital quantification of proteins and mrna in single cells. Nature communications, 10(1):1–10, 2019.

[37] Paul Macklin. When seeing isn’t believing: How math can guide our interpretation of measurements and experiments. Cell Systems, 5(2):92–94, 2017.

[38] Oliver J Maclaren and Ruanui Nicholson. What can be estimated? identifiability, estimability, causal inference and ill-posed inverse problems. arXiv preprint arXiv:1904.02826, 2019.

[39] Hirotaka Matsumoto, Hisanori Kiryu, Chikara Furusawa, Minoru SH Ko, Shigeru BH Ko, Norio Gouda, Tetsutaro Hayashi, and Itoshi Nikaido. Scode: an efficient regulatory network inference algorithm from single-cell rna-seq during differentiation. Bioinformatics, 33(15):2314–2321, 2017.

[40] Nicholas AM Monk. Oscillatory expression of hes1, p53, and nf-*κ*b driven by transcriptional time delays. Current Biology, 13(16):1409–1413, 2003.

[41] Matthew A Nunes and David J Balding. On optimal selection of summary statistics for approximate bayesian computation. Statistical Applications in Genetics & Molecular Biology, 9(1), 2010.

[42] Fabian Pedregosa, Gaël Varoquaux, Alexandre Gramfort, Vincent Michel, Bertrand Thirion, Olivier Grisel, Mathieu Blondel, Peter Prettenhofer, Ron Weiss, Vincent Dubourg, et al. Scikit-learn: Machine learning in python. the Journal of machine Learning research, 12:2825–2830, 2011.

[43] Dennis Prangle. Summary statistics in approximate bayesian computation. arXiv preprint arXiv:1512.05633, 2015.

[44] Christian P Robert, Jean-Marie Cornuet, Jean-Michel Marin, and Natesh S Pillai. Lack of confidence in approximate bayesian computation model choice. Proceedings of the National Academy of Sciences, 108(37):15112–15117, 2011.

[45] Markus Schirle, Marie-Anne Heurtier, and Bernhard Kuster. Profiling core proteomes of human cell lines by one-dimensional page and liquid chromatography-tandem mass spectrometry. Molecular & Cellular Proteomics, 2(12):1297–1305, 2003.

[46] Björn Schwanhäusser, Dorothea Busse, Na Li, Gunnar Dittmar, Johannes Schuchhardt, Jana Wolf, Wei Chen, and Matthias Selbach. Global quantification of mammalian gene expression control. Nature, 473(7347):337–342, 2011.

[47] Matthew J Simpson, Ruth E Baker, Sean T Vittadello, and Oliver J Maclaren. Practical parameter identifiability for spatio-temporal models of cell invasion. Journal of the Royal Society Interface, 17(164):20200055, 2020.

[48] Scott A Sisson, Yanan Fan, and Mark Beaumont. Handbook of approximate Bayesian computation. CRC Press, 2018.

[49] Stephen Smith and Ramon Grima. Spatial stochastic intracellular kinetics: A review of modelling approaches. Bulletin of mathematical biology, 81(8):2960–3009, 2019.

[50] Thomas R Sokolowski, Joris Paijmans, Laurens Bossen, Thomas Miedema, Martijn Wehrens, Nils B Becker, Kazunari Kaizu, Koichi Takahashi, Marileen Dogterom, and Pieter Rein Ten Wolde. egfrd in all dimensions. The Journal of chemical physics, 150(5):054108, 2019.

[51] Audrius B Stundzia and Charles J Lumsden. Stochastic simulation of coupled reaction–diffusion processes. Journal of computational physics, 127(1):196–207, 1996.

[52] Marc Sturrock, Andreas Hellander, Anastasios Matzavinos, and Mark AJ Chaplain. Spatial stochastic modelling of the hes1 gene regulatory network: intrinsic noise can explain heterogeneity in embryonic stem cell differentiation. Journal of The Royal Society Interface, 10(80):20120988, 2013.

[53] Marc Sturrock, Alan J Terry, Dimitris P Xirodimas, Alastair M Thompson, and Mark AJ Chaplain. Spatio-temporal modelling of the hes1 and p53-mdm2 intracellular signalling pathways. Journal of theoretical biology, 273(1):15–31, 2011.

[54] Mikael Sunnåker, Alberto Giovanni Busetto, Elina Numminen, Jukka Corander, Matthieu Foll, and Christophe Dessimoz. Approximate bayesian computation. PLoS Comput Biol, 9(1):e1002803, 2013.

[55] David J Warne, Ruth E Baker, and Matthew J Simpson. Using experimental data and information criteria to guide model selection for reaction–diffusion problems in mathematical biology. Bulletin of Mathematical Biology, 81(6):1760–1804, 2019.

[56] Alexandra S Whale, Jim F Huggett, Simon Cowen, Valerie Speirs, Jacqui Shaw, Stephen Ellison, Carole A Foy, and Daniel J Scott. Comparison of microfluidic digital pcr and conventional quantitative pcr for measuring copy number variation. Nucleic acids research, 40(11):e82–e82, 2012.

[57] Richard A Williams, Jon Timmis, and Eva E Qwarnstrom. Computational models of the nf-kb signalling pathway. Computation, 2(4):131–158, 2014.

[58] Samuel Wiqvist, Pierre-Alexandre Mattei, Umberto Picchini, and Jes Frellsen. Partially exchangeable networks and architectures for learning summary statistics in approximate bayesian computation. In International Conference on Machine Learning, pages 6798–6807, 2019.

